# DNA tetrahedron as a carrier of doxorubicin for metastatic breast cancer treatment

**DOI:** 10.1101/2023.08.02.551657

**Authors:** Payal Vaswani, Naveena A Hema, Krupa Kansara, Landon Dahle, Ashutosh Kumar, Dhiraj Bhatia

## Abstract

Metastatic breast cancer is a significant clinical challenge calling for novel and efficient therapeutic approaches. DNA tetrahedron, a highly programmable nanocage, offers some promising attributes including biocompatibility, stability, and functionalization making it an attractive candidate for drug delivery. In this study, we have explored the potential of DNA tetrahedron as a carrier of doxorubicin, a DNA and RNA synthesis-inhibiting chemotherapy drug. We have encapsulated doxorubicin in DNA tetrahedron (TD: Dox) and subsequently focused on metastatic breast cancer cells for the effect of the same. We showed that TD: Dox has the potential to inhibit the migration of cancerous cells in the 2D model and inhibit the invasion of tumor cells in the 3D model as well. This system also can be uptaken in *in vivo* zebrafish model as well. Overall, this study promises the TD: Dox system as an ideal drug delivery model and a viable approach for metastatic breast cancer treatment.

## 1. Introduction

Breast cancer is a widespread, potentially life-threatening ailment that continues to be a major global health concern. Worldwide, it is the most prevalent cancer in women, in developed and developing countries both. 2.3 million new cases and 0.7 million deaths were reported in 2020 for breast cancer by WHO. Breast cancer has been reported number one cancer in females in India^11^. The current treatments include surgery, chemotherapy, and radiation. One of the major problems in treatment is metastasis of cancer, which is the ability of cancerous cells to migrate to different body parts and form a secondary cancer^22^. Doxorubicin (Dox) is a widely used anti-cancer agent which primarily targets the synthesis of DNA and RNA^33^. However, there is the issue of low cellular intake and efflux via P-glycoprotein which poses an issue in Dox accumulation in cancerous cells^44^. This poses a requirement for a delivery system that can focus on delivering the Dox to cancerous cells.

DNA Nanotechnology is an active field focusing on the design and assembly of DNA molecules at the nanoscale to address the problems of the biological world. DNA being a versatile material, it has immense potential for programmability and creating different structures. It is biocompatible and non-immunogenic making it ideal for use in biological systems^55^. DNA tetrahedron (TD) is one of the most dynamic structures studied in this field^66^. TD is taken up by cells without the aid of transfection agents using endocytosis^77^. TD uptake is higher in cancerous cells compared to normal cells^88^ which can have substantial benefits as a drug delivery agent. The specificity and effectiveness of cancer treatments can be increased by using tailored medication delivery and therapeutics with potentially fewer off-target effects. Additionally, it has been demonstrated that TD influences the Wnt and Notch pathways, suggesting that it does influence signaling pathways^99^. This makes it the ideal choice for Dox delivery.

In this study, we aim to develop a drug delivery system by loading the chemotherapy drug Dox to TD. We studied the release kinetics of Dox from TD for its drug release potential. We primarily focused on understanding the cellular uptake of TD: Dox in MDA-MB-231 cells, metastatic breast cancer cells, and its effect on the migration and invasion ability of the cells. We also studied the potential of TD: Dox for DNA damage as Dox interferes with DNA and RNA synthesis. We further explored the uptake of TD: Dox in an *in vivo* zebrafish model. Overall, this study aims to pave the way for further research and translation of the TD: Dox model in clinical settings.

## 2. Results

### Synthesis and Characterization of TD and TD: Dox system

We synthesized TD using one pot synthesis method^1010^ (**Figure 1a**). Briefly, all four primers were mixed at equimolar ratio in reaction buffer containing 2 mM MgCl_2_ in nuclease-free water. They were subjected to thermal annealing from 95°C to 4°C. The final TD was further characterized using electrophoretic mobility shift assay (EMSA), dynamic light scattering (DLS), and atomic force microscopy (AFM). EMSA lies on the principle of band shift due to molecular weight change. We used 10% native PAGE to check the formation of higher-order structure and found the retardation of bands according to the stoichiometric ratio of the oligos added (**Figure 1b**). We carried out DLS next, to identify the hydrodynamic size of the TD and found out that the hydrodynamic radius is 11.7 ± 1.0 nm (**Figure 1c**). Morphology of TD was observed using AFM which showed the formation of triangular structures with approximately 20 nm size (**Figure 1d**).

**Figure 1.**
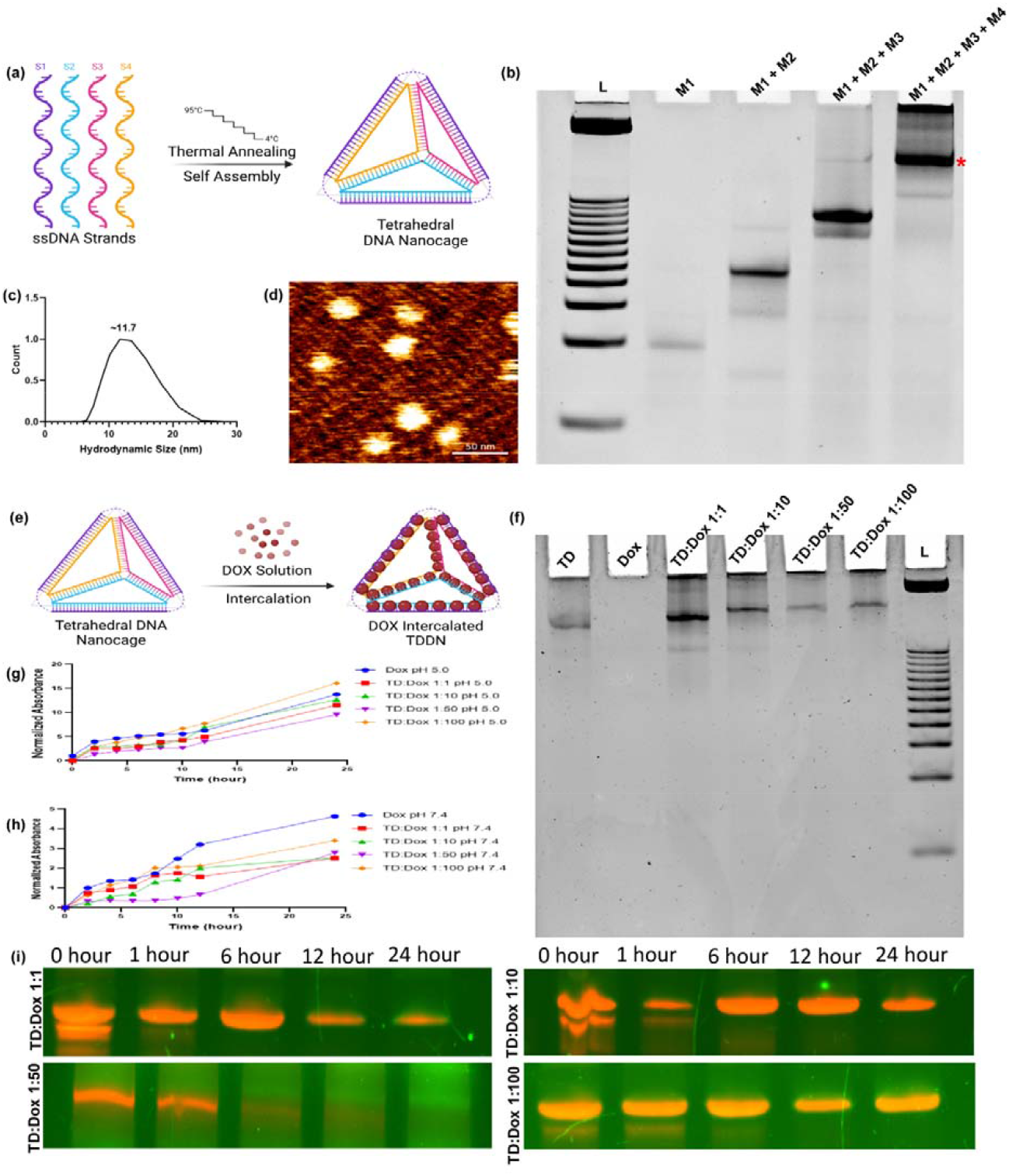
**(a)** Representative image for synthesis of TD **(b)** 10% Native PAGE showing synthesis of TD; lane 1 – Ladder, 2- M1, 3- M1+M2, 4- M1+M2+M3, 4-M1+M2+M3+M4 (TD) **(c)** Dynamic light scattering indicating the hydrodynamic radius of TD ∼ 11.7 nm **(d)** Atomic force microscopy showing morphology of tetrahedral particles of ∼ 20 nm size **(e)** Representative image for loading doxorubicin on TD **(f)** 10% Native PAGE showing TD loaded with doxorubicin in different concentrations; Lane 1- TD, 2- Dox, 3- TD: Dox 1:1, 4- TD: Dox 1:10, 5-TD: Dox 1:50,6- TD: Dox 1:100, 7-Ladder **(g)** Drug release study of free doxorubicin and TD loaded with doxorubicin with different concentrations at pH 5.0 **(h)** Drug release study of free doxorubicin and TD loaded with doxorubicin with different concentrations at pH 7.4 **(i)** Stability of TD: Dox system with 10% FBS at 37°C at 0, 1, 6, 12 and 24 hours shows robust stability of the cage with drug for upto 24 hours.

Once TD was synthesized, we moved to load doxorubicin on the TD (**Figure 1e)**. We loaded different molar ratios of doxorubicin (1, 10, 50, and 100 μM) to a constant ratio of TD (1 μM). Loading of the drug occurred at 37°C for 3 hours with shaking at 350 rpm and TD (1 uM) and Dox (100 uM) served as controls. After the reaction, unloaded Dox was removed by centrifugation at 10000 rpm for 10 minutes and the supernatant was collected^11^.^11^ The loaded cages were then desalted to remove any excess unbounded dox. To analyze the loading, EMSA was used. 10% native PAGE was run and there was a band shift as the amount of doxorubicin increased. Since doxorubicin and EtBr intercalates with DNA, as the doxorubicin increased, the band intensity of EtBr-stained gel decreased which additionally confirmed the loading of doxorubicin with TD (**Figure 1f)**.

To determine whether Dox could be released from TD, we performed a drug release study. We used dialysis bags of 3.5 MWCO with pH 5.0 and pH 7.4 phosphate buffer systems. We found that at pH 5.0 and pH 7.4, the free drug release is faster compared to loaded cages (**Figure 1g and Figure 1h**). However, at pH 5.0, the TD: Dox 1:100 has more release at 24 hours compared to a free drug (**Figure 1g**). This might be because TD enters the cells via clathrin-mediated pathway^88^ and endosomal pH is near 5.0^1212^ helping the release of drug from the TD.

For using these nanostructures in cellular and biological applications, the stability of nanostructures is a key factor as they are extremely vulnerable to nuclease destruction. The stability of the TD: Dox system was checked with 10% FBS at 37°C, and it showed that loaded structures are very stable till 24 hours as well (**Figure 1i**). This might be due to the intercalation of Dox may reduce the amount of freely available nuclease action sites.

#### Cellular Uptake of TD: Dox

A previous study from our lab has shown that TD can enter the cells within 15 minutes^88^. We wanted to check whether TD: Dox system could enter the cells or not. We incubated the MDA-MB-231 cells, which are metastatic breast cancer cells, with TD, Dox, and TD: Dox systems for 1 hour. Dox is a fluorescent drug that came into advantage for quantifying the amount of drug entering the cell. We used confocal laser scanning microscopy to study quantitatively and qualitatively the uptake of TD: Dox systems. We found out that with an increase in the concentration of Dox loaded in TD, the fluorescence intensity of Dox increased (**Figures 2a and 2b**). Interestingly, the intensity of Dox in TD: Dox 1:100 is more compared to free Dox in 60 minutes. The mechanism of action of Dox is to intercalate to DNA and disrupt the DNA replication^13^.^13^ To compare the binding of Dox to the DNA, we quantified the fluorescence intensity of DAPI as well (**Figure 2c**), which binds to DNA. We found that as the amount of Dox increases, the fluorescence of DAPI decreases. This shows that doxorubicin has been able to reach the nucleus and interact with the DNA confirming the release of Dox from the TD: Dox system. The confocal cell images also show an accumulation of Dox in the nucleus.

**Figure 2.**
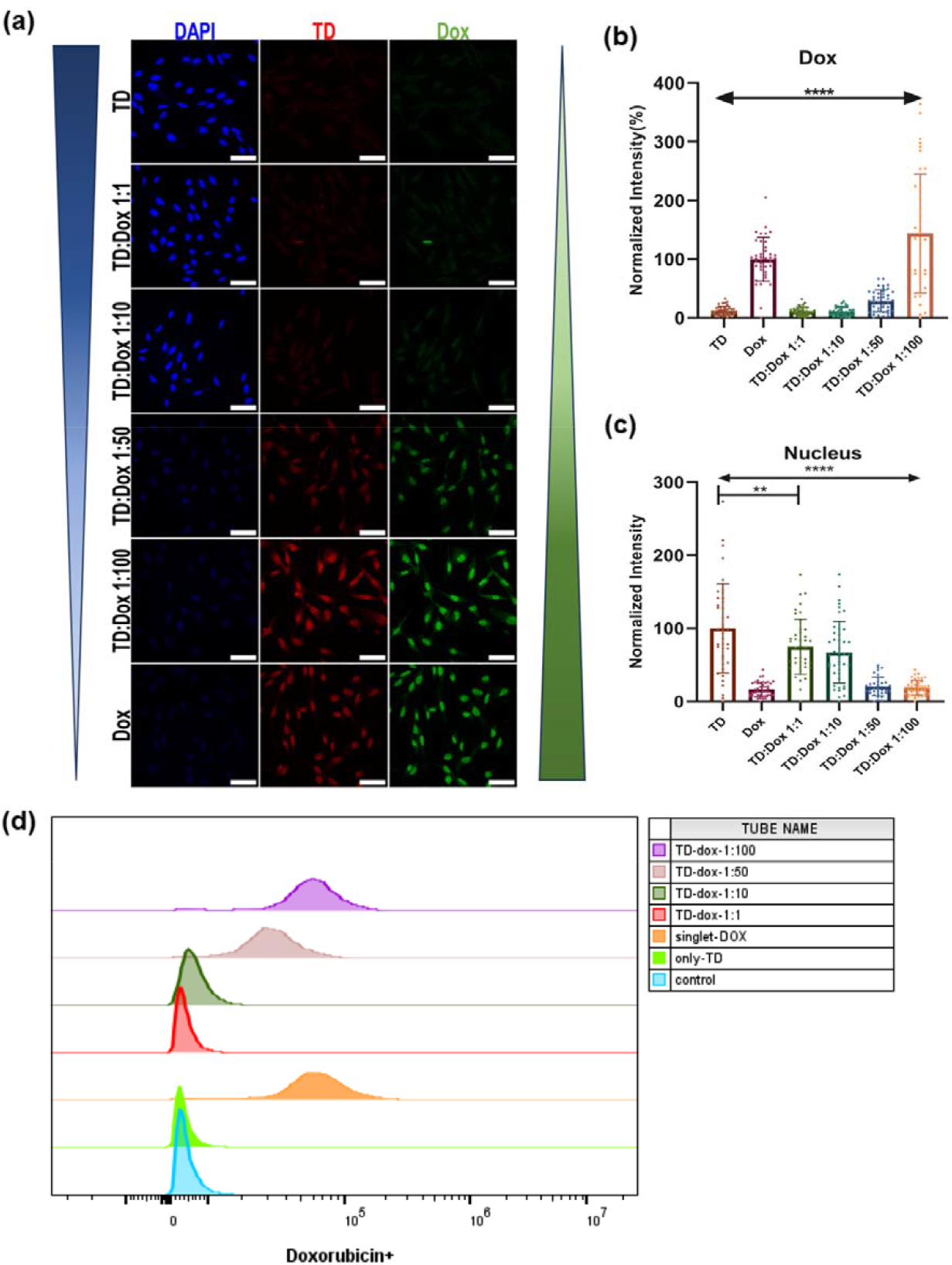
**(a)** Cellular Uptake of TD: Dox system in MDA-MB-231 cells for 60 mins. Scale bar is 20 μM **(b)** Quantification of Doxorubicin accumulated in cells with TD: Dox system for 60 mins **(c)** Quantification of DAPI accumulated in cells with TD: Dox system for 60 mins **(d)** Doxorubicin positive population gated using flow cytometry. ** indicates p <0.01, **** indicates p<0.0001

To further confirm the uptake of Dox, we performed flow cytometry. The experimental conditions were the same and uptake for 1 hour was checked. We acquired 10000 events and gated them further for Dox positive population. The flow cytometry results aligned with the confocal results and showed that TD: Dox 1:100 has more amount of Dox accumulation inside the cells compared to free Dox. Moreover, we can also see the increasing level of Dox accumulation with an increase in the amount of Dox loaded in the TD (**Figure 2d**).

### Migration and Invasion Potential

Since MDA-MB-231 is a metastatic breast cancer cell line, we wanted to know if TD: Dox system can inhibit the cell migration in turn decreasing the chances of metastasis. A previous study by Gada et al. has shown that TD can promote migration^14^.^14^ So, we performed scratch assay to understand that after loading Dox, whether the migration potential of cells increases or decreases. Control and TD showed good migration capacity of the cells (**Figure 3a, 3b**). While Dox inhibited the migration of the cells in 24 hours and the cell morphology drastically changed indicating the inherent property of Dox to damage the cells. For the TD: Dox systems, a 1:1 ratio showed similar results as TD as the drug concentration on TD is not enough to suppress the property of TD to help in migration. But as the Dox concentration increased, the migration of cells decreased. At 24 hours, the cell morphology also changed in TD: Dox 1:50 and TD: Dox 1:100 which was similar to free Dox indicating that these concentrations are more effective compared to TD: Dox 1:1 and TD: Dox 1:10.

**Figure 3.**
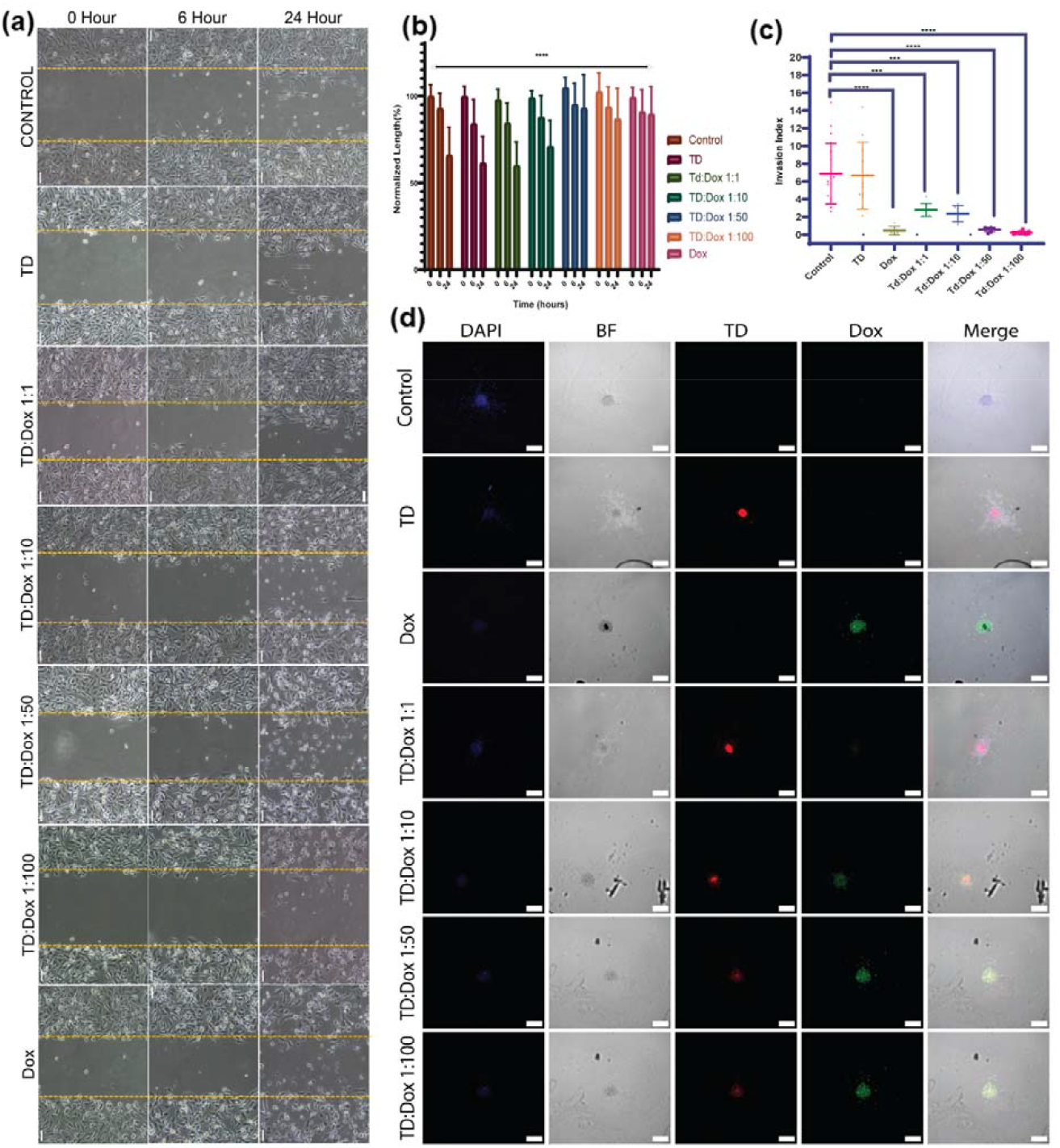
**(a)** Migration study of MDA-MB-231 cells with TD: Dox system. Scale bar is 200 μM **(b)** Quantification of wound distance with TD: Dox system **(c)** Quantification of invasion index for spheroids treated with TD: Dox system. Scale bar is 100 μM **(d)** Invasion study of MDA-MB-231 spheroids with TD: Dox system. n= 3 independent experiment; ** indicates p <0.01, *** indicates p<0.001, **** indicates p<0.0001

We then checked the invasion index in the 3D spheroid model as it is a more accurate representation of the tumor compared to the 2D cell culture. We prepared spheroids using the hanging drop method. The formation of spheroids was checked using bright field microscopy and then transferred on coverslips with a 3:1 ratio of collagen: media and incubated for 30 mins at 37°C. They were treated with TD: Dox systems for 24 hours and subsequently fixed and mounted. They were then analyzed with confocal microscopy at 10x magnification. The results showed a pattern similar to migration assay (**Figure 3c, 3d**). Control and TD showed a similar amount of invasion index while Dox significantly inhibited the invasion of spheroids compared to control. With the TD: Dox systems, as the concentration of Dox increased, the invasion index decreased. The TD: Dox 1:100 system showed an invasion index a little lower than free Dox. This indicates that TD: Dox 1:100 system can help in inhibiting the metastasis of the tumor.

#### DNA Damage and In Vivo Study

We further set out to determine whether Dox impacts cells and nuclei upon being uptaken in cells. Dox has been found to damage DNA^1515^, therefore we decided to conduct a comet assay to check for DNA damage^1616^. We used alkaline comet assay and 0.75% low melting agar to mix with the cells and formed a thin layer over pre-gelled slides. We scored the comets and used olive tail movement to quantify the level of DNA damage. Since TD: Dox 1:100 system showed better results compared to free dox, we used that alone for further experiments. The result showed significantly higher DNA damage in TD: Dox 1:100 system compared to control (**Figure 4a, 4b**). This shows that Dox is released from TD: Dox 1:100 system, reaching the nucleus and damaging the DNA.

**Figure 4.**
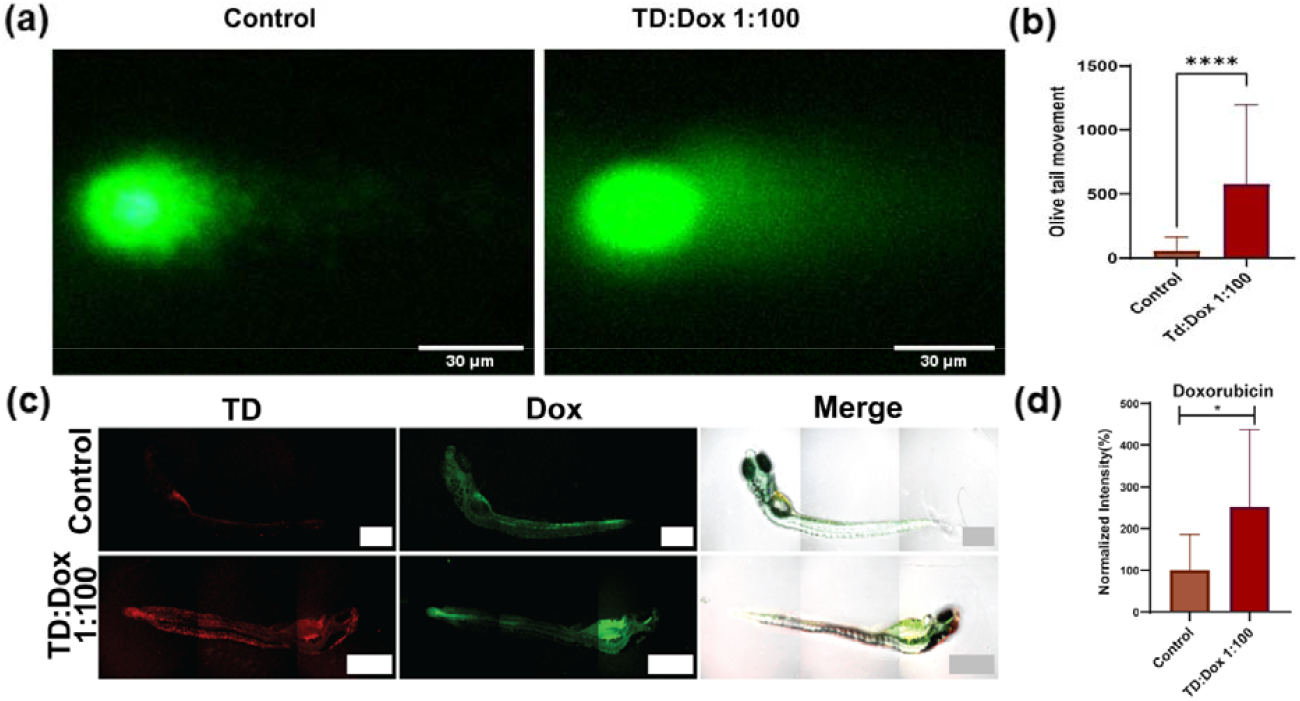
**(a)** Comet assay representative image showing control with very little tail and TD: Dox 1:100 with a long tail **(b)** Quantification of olive tail moment for comet assay (graph represents median with interquartile range) **(c)** Quantification of doxorubicin intensity in zebrafish model **(d)** Quantification of tetrahedron intensity in zebrafish model **(e)** Uptake of TD: Dox 1:100 in in vivo zebrafish model. * indicates p <0.05, **** indicates p<0.0001

We also sought to investigate the behavior of the TD: Dox system in an animal model. Using a zebrafish model, we evaluated the TD: Dox system. The results suggest that the TD: Dox system can enter zebrafish within 4 hours and doxorubicin gets accumulated at various regions of the fish (**Figure 4c, 4d, 4e**). This promises the TD: Dox system for in vivo delivery of Dox.

## 3. Discussions and Conclusions

Our goal was to use TD as a carrier of Dox for drug delivery in treating metastatic breast cancer. To load the drug, we utilized intercalating property of Dox in double-stranded DNA. The π-π stacking will aid in loading more Dox with rising concentration as we observed in the case of TD: Dox 1:100 system^17^. When we tested for drug release, the results were comparable to those of Xia et al., who reported that the drug release increased at pH 5^18^. TD enters the cells via clathrin-dependent endocytosis, where it will enter the endosomes^8,19^. The endosomal pH is acidic and will be favorable for the release of Dox. We also discovered that TD: Dox system’s stability increased as Dox concentration increased, which is a big step toward using this system for clinical models. Xu. et. al. showed that the inside corner of TD is more favorable for drug loading^20^. TD entry into the cells is also facilitated via a corner angle-mediated molecular interaction^21^. This should assist the entry of TD: Dox into the cells and release the drug. Zhang et. al. showed that TD: Dox system can enter the cells which are similar to our system^22^. We also showed that Dox can accumulate in the nucleus which is analogous to the study by Liu et. al.^23^. Our study showed the potential for TD: Dox system to inhibit cancer cell migration and invasion which is unique from the other studies. Future research on the long-term release, clearance, and safety of the TD: Dox system would be crucial for translating this model into clinical settings.

To conclude, the use of 3D DNA nanocage system to carry doxorubicin drug in the treatment of metastatic breast cancer cells shows tremendous promise. TD: Dox system showed enhanced cellular uptake in metastatic cells compared to free doxorubicin. We showed that the TD: Dox system can inhibit migration and invasion of cancerous cells making them valuable for the treatment of metastatic cells. The TD: Dox system can enter zebrafish without any additional agents making it a promising tool for in vivo models as well. However, additional preclinical and clinical investigations will help in optimizing the safety, dose, scalability, and long-term efficacy.

## 4. Materials & Methods

### 4.1. Materials

The primers (M1-M4) were obtained from Sigma Aldrich and the sequence is mentioned in Table 1. Nuclease-free water and magnesium chloride are obtained from SRL, India. Acrylamide: Bisacrylamide (30%), TEMED, APS, Penstrap, paraformaldehyde, ethidium bromide, triton X is obtained from Himedia. Doxorubicin hydrochloride, 6X loading dye, 25 bp ladder, and mowiol were obtained from Sigma Aldrich. The desalting column was obtained from Zeba. Dialysis membrane (3.5 kDA MW) is obtained from Thermo Fischer. DMEM Media, FBS, and trypsin are obtained from Gibco. Collagen was purchased by Corning.

**Table 1.**
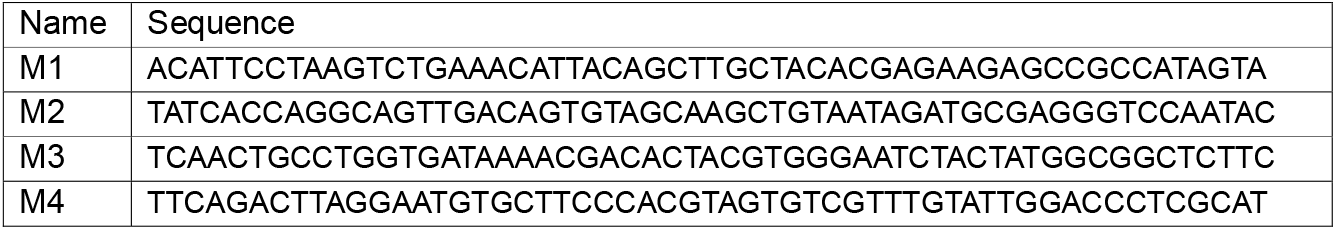

### 4.2. Synthesis and Characterization

#### Synthesis of TD

TD was synthesized using one-pot synthesis. Briefly, the four oligonucleotides were mixed in equimolar ratios (M1:M2:M3:M4 1:1:1:1) with the addition of 2 mM MgCl2. The reaction was heated to 95 °C before being gradually annealed and cooled till 4 °C. The reaction cycle comprised a 5 °C step decrease with a 15-minute interval between each step. The final concentration of TD formed was 2.5 μM.

#### Synthesis of TD: Dox

TD: Dox system was prepared with 4 concentration variations. TD concentration was kept constant at 1 μM and Dox concentration was 1 μM, 10 μM, 50 μM and 100 μM for TD: Dox 1:1, TD: Dox 1:10, TD: Dox 1:50, and TD: Dox 1:100 respectively.

#### Characterization of TD and TD: Dox

##### Electrophoretic Mobility. Shift Assay (EMSA)

EMSA was performed using 10% Native-PAGE to confirm the formation of a higher-order structure. The sample contained 5 μL of TD, 3 μL of 1× TAE loading buffer, and 1.5 μL of 6× loading dye. The gel was run at 100 V for 60 min. The gel was stained with EtBr stain and visualized using a Gel Documentation system (Biorad ChemiDoc MP Imaging System).

##### Dynamic Light Scattering (DLS)

The size-based characterization of TD was done using DLS to measure its hydrodynamic size. 50 μL of 125 nM sample was used for analyzing the hydrodynamic radius using the Malvern analytical Zetasizer Nano ZS instrument. The readings were taken in triplicates, with 13 readings in one run.

##### Atomic Force Microscopy (AFM)

The morphology-based characterization was performed using Atomic Force Microscopy. A freshly cleaved mica was used. 10 μL of 1:10 diluted TD was overlaid on top of the mica sheet and allowed to dry in a desiccator. It was then imaged using BIO AFM (Bruker) in tapping mode and the image was processed using JPK software.

#### In Vitro Studies

##### Drug Release Studies

Dialysis: 400 μL of the sample was loaded in 3.5K MWCO dialysis tubing and kept for stirring at 350 rpm in pH 5.0 and pH 7.4 phosphate buffer at 37°C. Samples of the media were taken every 2 hours for 24 hours to test for the level of DOX released into buffers over time.

### 4.3. Cell Culture

MDA-MB-231 cells were maintained in DMEM media supplemented with 10% FBS and 1% PenStrap at 37°C and 90% humidity.

#### Cellular Uptake Using Confocal Microscopy

MDA-MB-231 cells were seeded in 24 well plates on coverslips and allowed to adhere for 24 hours. The experiment was done once the cells were around 80% confluency. The media was decanted, and cells were washed with 1X PBS. The cells were serum starved for 20 mins. The cells were then treated with 200 nM systems (TD, TD: Dox 1:1, TD: Dox 1:10, TD: Dox 1:50, TD: Dox 1:100) or 20 uM Dox for 1 hour in serum-free media at 37°C. The media was removed, and cells were washed with 1X PBS two times to remove unbound systems. The cells were further fixed with 4% PFA for 15 mins at 37°C. They were washed with 1X PBS three times to remove any PFA crystals. They were mounted on slides with Mowiol + DAPI and stored at 4°C for drying. The uptake was then analyzed with Leica SP8 Laser Scanning Confocal Microscope using 405 nm, 488 nm, and 633 nm lasers and 512*512 resolution using 63X oil immersion objective. The data was further processed using Fiji ImageJ software.

#### Flow Cytometry

The cells were grown in 6 well plates and allowed to adhere for 24 hours. The experiment was done once the cells were around 80% confluency. The media was decanted, and cells were washed with 1X PBS. The cells were serum starved for 20 mins. The cells were then treated with 200 nM systems (TD, TD: Dox 1:1, TD: Dox 1:10, TD: Dox 1:50, TD: Dox 1:100) or 20 uM Dox for 1 hour in serum-free media at 37°C. The media was removed, and cells were washed with 1X PBS two times to remove unbounded systems. The cells were then trypsinized and the pellet was collected, and further dissolved in filtered 1X PBS. They were immediately subjected to BD FACS Aria Fusion to analyze the cellular uptake. The data was further processed using FlowJo software.

#### Scratch Assay

The cells were seeded in a 6-well plate and allowed to grow till 100% confluency. Scratch was created at the center of the wells using a sterilized 200 μL tip. The wells were then gently washed with 1X PBS to remove the detached cells. They were incubated with fresh serum-free media with 200 nM of DNA cage systems (TD, TD: Dox 1:1, TD: Dox 1:10, TD: Dox 1:50, TD: Dox 1:100) or 20 μM Dox, and control as untreated cells. They were imaged at different time points (0, 6, 24 hours) at 10X in the Nikon microscope. The images were further processed using Fiji Image J software. The distance between the scratch was measured to access the wound closure.

#### 3D Invasion assay using spheroids

MDA-MB-231 cells were used to prepare 3D spheroids using the hanging-drop method. The cells were trypsinized, 100 cells were taken per 50 μL of complete media and seeded in droplet form on the lid of a Petri dish. The base of the Petri dish was filled with 15 mL of autoclaved water to provide humidity for spheroid growth. The cells were incubated at 37°C for 24 h. The cells clumped together and aggregated due to gravity to form spheroids. The spheroid formation was confirmed by observation under a bright field microscope. The spheroids were transferred to 24 well plates using 3:1 collagen to media proportion on coverslips, and it was incubated at 37 °C for 30 minutes. The spheroids were then incubated with 200 nM of different cage systems (TD, TD: Dox 1:1, TD: Dox 1:10, TD: Dox 1:50, TD: Dox 1:100) or 20 uM Dox and untreated spheroids were control in complete media for 24 h at 37°C. The media was decanted, and spheroids were washed once with 1X PBS. The spheroids were fixed using 4% PFA for 20 min at 37°C. Spheroids were washed gently twice with 1X PBS and mounted using Mowiol+DAPI.

#### Comet Assay

The treatment protocol for the assay remains the same as cellular uptake. Briefly, 50 μL of cells were mixed with 100 μL of 0.75% low melting agar and placed on pre-treated slides (1% normal melting agarose), covered with coverslips, and allowed to gel on a cold plate for 5 min. Next, coverslips were carefully removed and one more layer of low-melting agar was added and allowed to solidify. The slides were then placed in lysing solution at 4°C for 2 hours. They were then placed in an electrophoresis chamber with electrophoresis buffer with pH>13 and left for 20 mins for unwinding. The electrophoresis was performed for 25 mins at 300 mA current. The slides were then washed with neuralization buffer 3 times for 5 minutes. They were stained with SyBr Gold and the image was acquired using a 10X objective of Nikon microscope. The images were further processed using Fiji ImageJ software with the Opencomet plugin.

### 4.4. *In vivo* uptake in zebrafish embryos

The uptake assays of Td: Dox were conducted following the guidelines of the Organization for Economic Co-operation and Development. The experiments were performed on embryos 9 days after hatching. Any dead embryos were removed, and the remaining ones were placed in 6-well plates, with 15 eleuthero embryos in each well. A group of eleuthero embryos was treated with Td: Dox (1:100) at the concentration of 200nM, and another group served as control, receiving no nanoparticles. After a 4-hour incubation with the nanoparticles, the medium was replaced with fresh E3 media to remove any excess nanoparticles. Then, the eleuthero embryos were washed and fixed with a 4%PFA for 2 minutes. Following fixation, the embryos were mounted along with Mowiol and allowed to dry before further analysis using confocal imaging.

### 4.5. Statistical Analysis

The statistical analysis was performed using GraphPad Prism 8.0. The data presented are mean with SD unless mentioned otherwise. The test used was the t-test and one-way ANOVA for normally distributed data and the Mann-Whitney test for non-parametric data. All the experiments were performed in triplicates. The statistical significance is denoted by * is p<0.05, ** is p<0.01, *** is p<0.001 and **** is p<0.0001.

## Acknowledgments

All authors thank IITGN and CIF-IITGN for their facilities. PV thanks UGC for the Ph.D. fellowship and IITGN for the additional fellowship. HN thanks PMRF for the Ph.D. fellowship. KK thanks SERB for the NPDF fellowship. DB thanks SERB-DST for the Ramanujan fellowship, IITGN, DST, Gujcost-DST, and GSBTM for research grants. DB is a member of the Indian National Young Academy of Sciences, INYAS-INSA. The central instrumentation facilities at IITGN are acknowledged.

